# Shallow MinION sequencing to assist *de novo* assembly of the *Streptococcus agalactiae* genome

**DOI:** 10.1101/485029

**Authors:** Tamara Hernandez-Beeftink, Hector Rodriguez-Perez, Ana Díaz-de Usera, Rafaela Gonzalez-Montelongo, José M. Lorenzo-Salazar, Fabián Lorenzo-Díaz, Carlos Flores

**Affiliations:** Research Unit, Hospital Universitario Ntra. Sra. de Candelaria, Universidad de La Laguna, Santa Cruz de Tenerife, Spain; Genomics Division, Instituto Tecnológico y de Energías Renovables (ITER), Santa Cruz de Tenerife, Spain; Departamento de Bioquímica, Microbiología, Biología Celular y Genética, Universidad de La Laguna, Santa Cruz de Tenerife, Spain; CIBER de Enfermedades Respiratorias, Instituto de Salud Carlos III, Madrid, Spain

**Keywords:** Hybrid assembly, Nanopore, artefactual reads

## Abstract

Despite the reduced read length, the so-called Next-Generation Sequencing (NGS) of second-generation has allowed rapid and complete genome characterization of many species. MinION (Oxford Nanopore Technologies), a portable third-generation NGS device, enables sequencing of long DNA fragments at low cost. Here we used a low-coverage MinION sequencing in combination with short-read NGS to improve genome assembly. We tested this possibility by using MinION R9.0 with Rapid 1D kit and MiSeq with >300X paired-end 300 bp reads (Illumina, Inc.) for the genome assembly of a *Streptococcus agalactiae* clinical isolate (2.2 Mb). With as few as 1,171 MinION reads that covered the genome at 2.4X (the longest read being 186 Kb long), the hybrid assembly combining MinION and Illumina reads increased the N50 by 4.9-fold compared to the assembly using Illumina data alone. Almost 50% of the genome was represented into a single contig (1.02 Mb). Besides, this allowed the full reconstruction of mobile elements, including a plasmid, and improved gene annotation. Taken together, our results support that shallow MinION sequencing combined with high-throughput second-generation NGS constitutes a cost-efficient strategy for the assembly of whole genomes.

## Background

Next-Generation Sequencing (NGS) technologies have allowed sequencing and analysis Group B *Streptococcus* (GBS) genomes from different environments and organisms[1–6]. The presence of prophages and mobile elements in GBS was associated with the bacterial adaptation and ability to cause infections, likely enhancing their diversity and spread[7–10]. In fact, the estimates suggest that roughly 50% of *Streptococcus agalactiae* genome is grouped into islands of pathogenicity, where virulence genes and mobile elements are present[11]. *S. agalactiae* genome shows a complex organization and rearrangements, where the presence of a conserved backbone and 69 variable regions have been observed[12]. These observations justify the necessity of using *de novo* assembled genomes for better comparative bacterial studies and adequate tracking of mobile elements. However, the assembly of bacterial genomes based on short-read NGS technologies (so called second-generation) usually produce discontinuous sequences[13,14], often requiring the assistance from algorithms, alternative library approaches or sequencing technologies to provide sequence continuity.

Third-generation NGS technologies allow obtaining longer DNA reads from PCR-free DNA libraries in real time, constituting a solution for resolving repeats[15] and obtaining complete assembled genomes. In this context, the portability and reduced costs of sequencing with the MinION device (Oxford Nanopore Technologies) has facilitated the adoption of this technology in many distinct applications[16]. Besides its utility for the analysis of structural variants[17,18], SNP determination[19], cytosine methylation[20], transcriptomics[21], and in-field experiments in extreme environments[22], MinION has widely used to sequence small whole genomes from virus (10.8 kb)[23], mitochondrial DNA (16 kb)[24], and bacteria (4.6 Mb)[25], allowing their assembly into single contigs when depth of coverages were above 20–40X[26]. Nevertheless, this technology continues to be challenged by its high error rate (>10%)[27], particularly in homopolymeric regions[28], and the generation of reads that fail to align against the target[29,30]. Because of this, hybrid assemblies leveraging the per-base quality of high-coverage (>200X) second-generation data with the bridging capacity of >10X depth-of-coverage third-generation data is nowadays perceived as the optimal choice for obtaining accurate assemblies[24,27,31–35]. Given the large variability in sequence throughput offered by this technology well below product specifications, here we tested the benefit of shallow MinION sequencing data in assisting the hybrid genome assembly of a *S. agalactiae* clinical isolate (genome size of 2.2 Mb).

## Methods

### Bacterial isolate, growing conditions and validation

We have previously examined the IMESag-rpsI mobile element distribution in 240 whole genomes of Group B *Streptococcus* isolates. The *S. agalactiae* HRC strain was subjected to a deeper examination to assist the characterization of the mobile element in that study[36]. *S. agalactiae* HRC is a serotype V representative strain belonging to the tetracycline-resistance CC1 lineage Tn916–1[37], which was isolated in 2009 from a 60-year-old woman with abdominal sepsis at the Hospital Ramón y Cajal (Madrid, Spain)[36].

The bacterial isolate was cultured in TSB (Tryptic Soy Broth, Becton Dickinson) liquid medium at 37ºC for 24 h. DNA extraction was performed with the GenElute Bacterial Genomic DNA kit (Sigma-Aldrich) and was quantified on a Qubit 3.0 fluorometer using the dsDNA BR assay kit (Thermo Fisher Scientific). Its identity was first validated by 16S rRNA gene sequencing as follows. DNA was amplified by PCR with 30 cycles of 95°C for 20 s, 50°C for 30 s, and 72°C for 15 s, using the HotStarTaq master mix kit (QIAGEN), and the primers 1391R (5'-GACGGGCGGTGWGTRCA-3') and 27F (5'-AGRGTTYGATYMTGGCTCAG-3'). Amplified material was purified with ExoSAP-IT (Thermo Fisher Scientific), subject to Sanger sequencing with BigDye Terminator v3.1 Cycle Sequencing kit (Thermo Fisher Scientific), and products purified with DyeEx 2.0 Spin kit (QIAGEN), following the manufacturer's recommendations. Sequencing products were resolved and basecalled in a 3500 Genetic Analyzer (Thermo Fisher Scientific). A simple BLAST[38] search on the sequences obtained verified that the bacterial isolate had a 99% identity to *S. agalactiae*.

### Illumina sequencing

DNA from *S. agalactiae* HRC (1 ng) was used to generate sequencing libraries with the Nextera XT kit (Illumina, Inc.) following the manufacturer's recommendations. A MiSeq instrument and a MiSeq Reagent kit V3 (Illumina, Inc.) was used for 300 base paired-end sequencing along with a spike-in of 20% of the PhiX Control v3 (Illumina, Inc.). Library construction and the sequencing experiment were performed at the Instituto Tecnológico y de Energías Renovables (ITER).

### MinION sequencing

Rapid Sequencing kit (SQK-RAD001) (Oxford Nanopore Technologies) was used for library preparation starting from 200 ng of *S. agalactiae* HRC purified DNA. Sequencing with MinION used SpotOn Flow Cell Mk I R9 Version (Oxford Nanopore Technologies) and followed manufacturer's recommendations, except that the experiment was left running only for 22 h.

### Bioinformatics and statistical procedures

MiSeq Reporter Software v1.18.54 (Illumina, Inc.) was used to convert intensities to basecalls and generate the FASTQ file. MinION basecalling was performed with MinKNOW GUI 1.1.21 software (Oxford Nanopore Technologies) and all records were processed with poRe v0.21[39] to extract the obtained reads in FASTQ file format. Unicycler v0.4.1[40] was used with default parameters for hybrid *de novo* assembly of *S. agalactiae* HRC genome, combining Illumina and Nanopore datasets, which involved multiple rounds of short-read polishing. As a reference for comparisons, Unicycler was also used to assemble the Illumina data alone. Assembly comparisons were assessed with QUAST v4.0[41] using the *S. agalactiae* SS1 genome (NZ_CP010867.1) as the reference. The assembled contigs were visualized with Bandage v0.8.1[42]. Annotation of genes and elements in the contigs was done with IonGAP[43]. Mauve v2.4.0[44] was used to identify the insertion sites of the *S. agalactiae* HRC mobile elements in the assembly against the reference (NZ_CP010867.1).

In order to further explore raw MinION reads, per-read GC content distribution was first assessed to identify outliers (>3 SD from the mean). The existence of artefactual reads in the output[29] was evaluated through comparisons against the reconstructed *S. agalactiae* HRC genome, including the two characteristic elements identified. BLAST v2.6.0[38] alignments were used to isolate unaligned artefactual reads. Bio.SeqUtils library of Biopython release 1.70 was then used to assess their differences in length (bp), GC content (GC%) and mean quality (Q) score against the reads aligning to the *S. agalactiae* HRC assembled genome. A non-parametric Mann-Whitney U-test was used to assess the significance in the comparisons. To test the possibility that particular pores were generating more artefacts than expected, as has been suggested elsewhere[29], unaligned reads were mapped back to the 512 nanopores of the Flow Cell and significance was tested assuming a Poisson distribution. Finally, to evaluate their impact in *de novo* assembly of *S. agalactiae* HRC genome, we then excluded all unaligned reads from the dataset, and used Unicycler under the same conditions to reassess the assemblies.

## Results

### Summary of sequencing results

Sequencing with Illumina resulted in 2,975,356 paired-end reads of 300 bp with Phred>30, which translates into 354X depth-of-coverage of *S. agalactiae* genome. In comparison, the MinION run proceeded with a total of just 276 active pores, providing 1,171 reads with a mean length of 4.5 kb and a Q score average of 10.8, and ranging from 3.8 to 25.4. While the largest proportion of reads accumulated in the 1.86–9.0 kb range (the N50 of the output was 9.2 kb) the output included reads as long as 186 kb (which represents almost 10% of the *S. agalactiae* HRC genome). The total throughput (5.2 Mb) theoretically translates into a 2.4X depth-of-coverage of the bacterial genome, which is well below product specifications, despite all initial quality controls of the platform were passed. Finally, the per-read GC content average of reads was 44.3% (ranging from 10.3% to 95.9%), noticeably larger than that of the reference sequence (35.5%; NZ_CP010867.1).

### Assembly improvements by using MinION data

The Illumina data assembly generated 23 contigs (>500 bp) with a N50 of 147,211 bp, and the largest contig being 525,288 bp long. The length of the assembly from the Illumina data alone was very close to the *S. agalactiae* reference, accounting for a genome size of 2,144,732 bp, a GC content of 35.3%, and an aligning length against the reference of 2,049,610 bp (97.9% of consensus identity) (Table 1). In comparison, although the hybrid assembly including also combining the Illumina and MinION reads did not allow to obtain a fully resolved bacterial genome, mostly because of a major unresolved structure containing repetitive elements, rRNA operon copies and tRNAs (Fig 1), it reduced the number of contigs to less than a half (8 with >500 bp), having a N50 of 714,293 bp and the largest contig being 1,018,988 bp long. The hybrid assembly generated a total length of 2,159,060 bp, with 2,062,825 bp aligning against the reference (98.5% of consensus identity) (Table 1). Both assemblies also evidenced a small plasmid (2,491 bp) as an independent replicon, which perfectly aligned with the *Streptococcus oralis tigurinus* 2426 plasmid pST2426 (ASXA01000016.1), along with two other mobile elements detected in previously studies[36] (Fig 1). In terms of gene annotation, the hybrid assembly also outperformed the Illumina only assembly (2,153 vs. 2,135 genes, respectively) when compared to the reference (2,195 genes) (Table 1). Taken together, the small amount of reads provided by MinION (0.04% of total reads obtained for the bacterial isolate) allowed to increase the N50 of the genome assembly by 4.9-fold, to cover almost 50% of the bacterial genome in a single contig, and to improve the annotation of predicted genes in the assembly.

**Fig 1.**
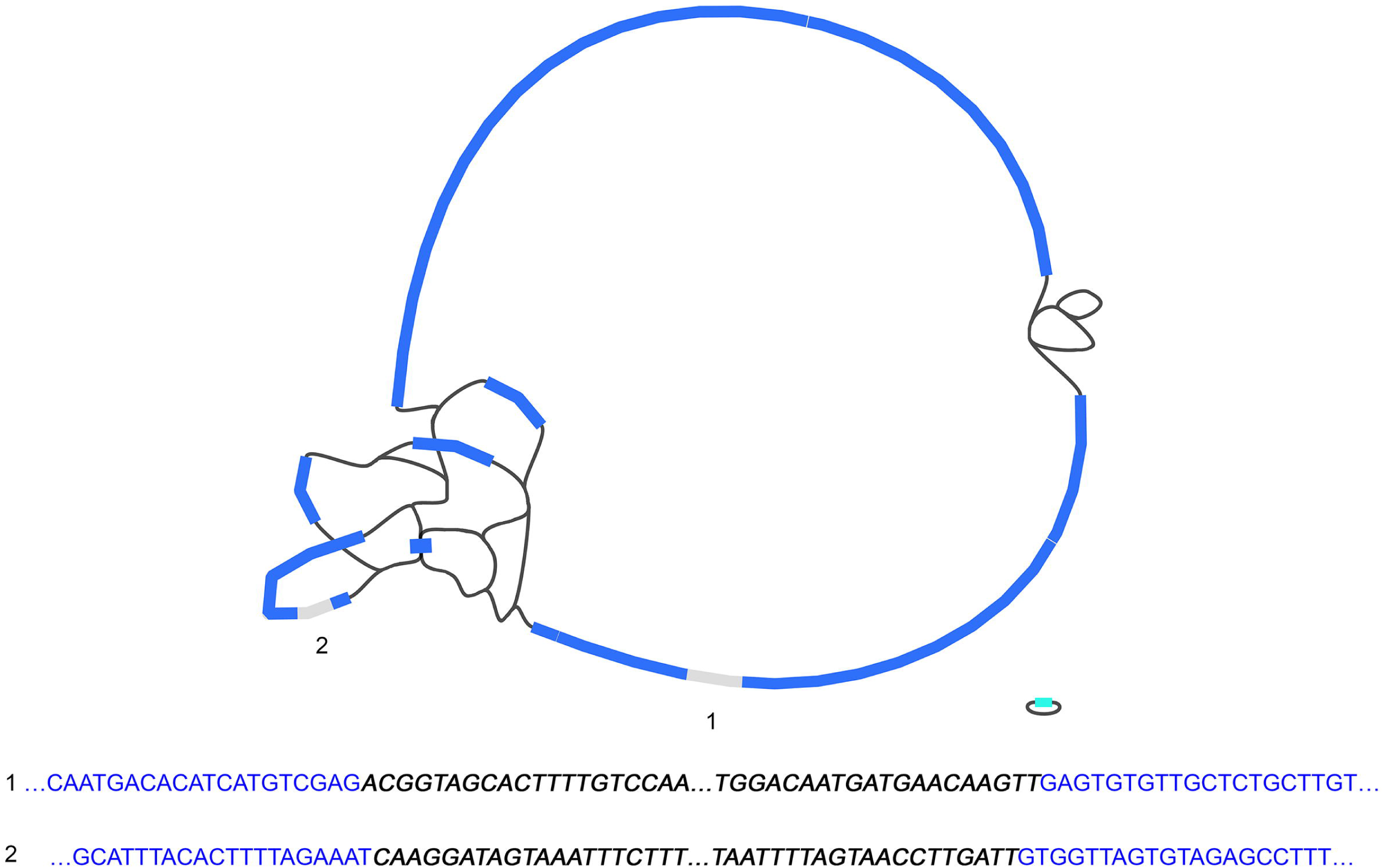
Schematic representation of the *S. agalactiae* HRC hybrid assembly. The flanking sequences of the two characteristics elements of this isolate are indicated at the bottom: 1) A conjugative element attached to the Tn916 transposon located in the largest contig (1,018,988 bp); 2) A prophage located in the third largest contig (224,154 bp). Most copies of the rRNA (16S, 23S, 5S) and tRNA genes were positioned within the largest unresolved assembly structure.

**Table 1.**
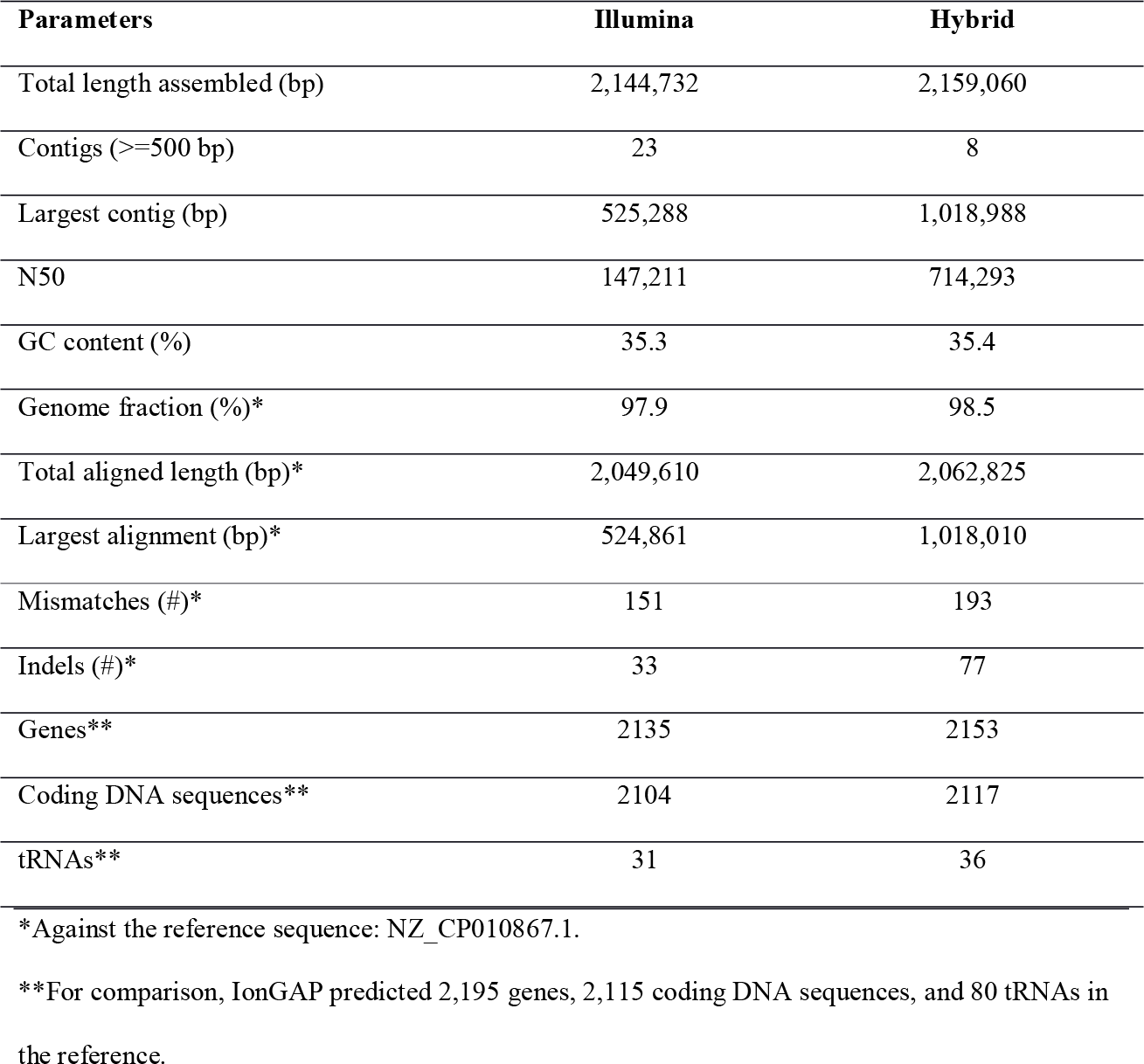
Summary of the hybrid and Illumina-only assemblies of *S. agalactiae* HRC genome.

### Characterization of unaligned MinION reads and impact on the assembly

A closer inspection of the MinION run records evidenced the existence of reads with large regions of limited sequence diversity (end to end) that were largely compatible with artifacts. Just by exploring the GC content of the MinION output, we detected 27 reads with large deviations (>3 SD from the mean) corresponding to GC contents below 7.2% or above 77.8%. The reads involved had a mean size of 716 bp, but were as large as 4.2 kb. However, in order to better identify and characterize the reads that were most likely artefactual, we used BLAST at 85% identity threshold to align all reads against the assembled genome. This allowed to identify 68 unaligned reads with a mean size of 2.0 kb (but as large as 21.6 kb) and a mean Q score of 6.7 (but as high as 22.9) (Fig 2). Only three of these unaligned reads (0.25% of total) aligned to sequences from other species or synthetic constructs according to a BLAST search (Table S1), most probably corresponding to experimental contaminants. Surprisingly, MinKNOW software only classified 20 of the 68 as failed based on internal manufacturer’s specifications. Unaligned reads differed significantly from reads that aligned for the GC content (74.2 vs. 42.5%; p= 2.5e^−37^), Q score (6.7 vs. 11.1; p= 2.2e^−21^), and mean read length (2.0 vs. 4.6 kb; p= 1.3e^−11^) (Fig 2). Strikingly, from the 276 active pores of the run, unaligned reads were obtained from 45, and two pores in particular experienced a significant accumulation of them (28.9%, p<0.019) (S1 Fig). Finally, a re-assessment of the hybrid assembly evidenced identical results with or without excluding unaligned reads, supporting that Unicycler was robust to their presence in the input. For simplicity, these results are not shown.

**Fig 2.**
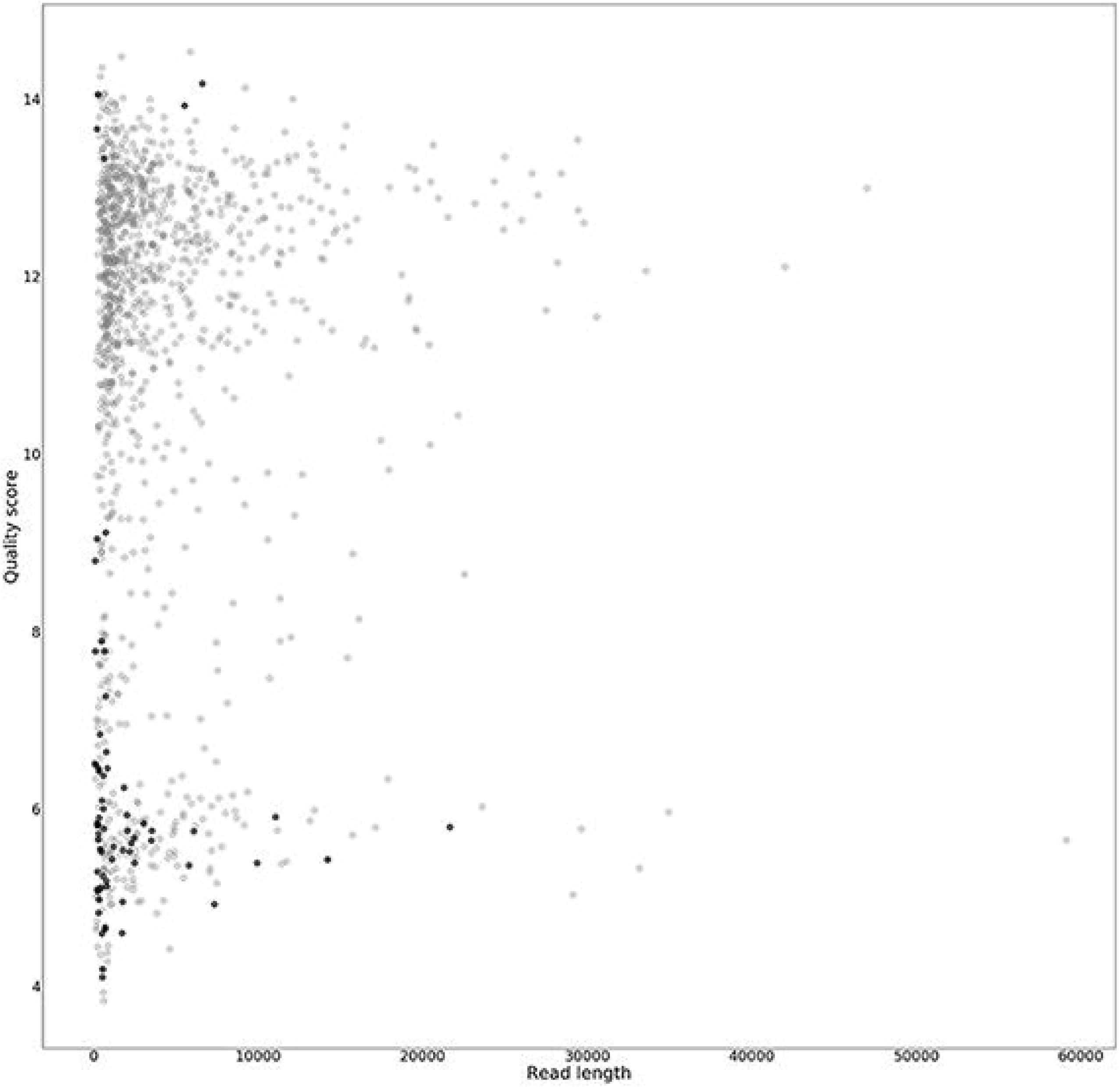
Distribution of MinION reads by quality (Q) scores and length (bp). Aligned reads are represented in grey, whereas the 68 unaligned reads are overlaid in black. For simplicity, the plot was capped to represent reads <60 kb length and Q scores <15.

## Discussion

Here we demonstrate that the use of very low amount of MinION reads combined with short-read sequencing data has the capacity to increase the continuity of assembly contigs and plasmid isolation without manual intervention, having the potential to fully resolve bacterial genomes. This was illustrated with the assembly of *S. agalactiae* HRC genome based on as few as 2.4X MinION depth-of-coverage data, paving the way for obtaining cost-effective continuum whole-genome sequences as the nanopore technology throughput increases. While other species may have alternative requirements, to the best of our knowledge, this is the first study assessing the benefits of using MinION reads to assist the complete assembly of microbial genomes with coverages well below 10X (usually assisted with >14X)[14,23,33,34]. Although we were unable to fully resolve the assembly, our results are quite reassuring considering the small number of MinION reads used in the experiment (0.04% of total reads), allowing to obtain almost the 50% of the bacterial genome in a single contig. Besides, we were able to detect and fully reconstruct a plasmid, previously attained only with higher depth-of-coverage and longer MinION reads[45].

High quality noise reads generated by MinION have been described recently in the literature[29], consisting on reads that did not map to the target, lab contaminants, or spike-in sequences. Many of those appeared to be artefacts of the technology that were associated with specific active pores[29], to which some authors have previously attributed an insufficient read quality[30]. In this study, MinION generated around 5% of reads of variable quality that did not map to the target. We found that despite a few of them had partial identity with bacterial sequences that may indicate minor experimental contaminations as has been shown by others[15,46], most of them consisted in low-complexity sequences from end to end. We demonstrate that a number of them were generated from specific active pores during our experiment. Based on these evidences, we speculate that most of these reads are artefacts produced during the sequencing process and that they are most likely independent from the basecaller being used. In this study, we demonstrate that they did not negatively affect the hybrid assembly process in the conditions assessed. However, their impact in other projected applications (such as shotgun metagenomics or whole-genome sequencing of eukaryotes) or algorithms other than Unicycler remain unknown. Solutions to reduce or identify and filter out these reads will need to be implemented.

With sufficient MinION yield providing enough depth-of-coverage, the standard analysis is based on long-read and error-tolerant assemblers such as Canu[47]. Because of the high error rate of this technology at the moment (10–20%)[27,46,48], and the problems resolving the homopolymers[28], the assembly process from MinION data alone typically necessitates of a number of error correction steps to produce high quality genome sequences[49]. Most importantly, this assembly process is computationally intensive[49,50]. Theoretically, MinION can produce many thousands of reads and a few dozen Gb per run[49]. However, in practice MinION outputs are highly variable translating sometimes in a few thousand reads (3,400 in Benitez-Paez et al[51]) and Mb scale outputs. In our hands, regular experiments are roughly in the scale of 100 Mb per run. This is because the number of active pores changes during the experiment (between 105 and 426 in Greninger et al[46]), below theoretical maximum in distinct runs, and the effective number of useful reads represents a fraction of the output[15,46]. Some studies have reported that a large proportion of the output (>50%) have failed to align against the target[30,52]. Besides, sudden crashes of the MinION software controller[14] or the computer connection failures[53] have been reported during a sequencing experiment. In addition, unofficial information provided by users in The Nanopore Community report extremely low occupancy of pores (<10%) during the sequencing experiments with Rapid 1D kits, which have been suggested to be related to the kit production itself. These numbers are comparable to our observations in this study and all these issues have obvious consequences in the throughput of a MinION experiment. Therefore, although not the standard, the limited yield obtained by the MinION sequencing in this study is not uncommon. In this context, a hybrid assembly with a very shallow coverage of MinION data constitutes an advantage over the approaches based on MinION data alone for obtaining cost-effective high-quality continuous bacterial genomes. Besides being computationally faster, this hybrid option reduces the burden of generating the thousands of long reads that are necessary to obtain high quality bacterial genome assemblies with MinION data alone (>20X)[26].

## Conclusions

We demonstrate that the combination of a small proportion of long reads with high-coverage short-read data constitutes a promising strategy to generate high quality, highly contiguous assemblies, here allowing to nearly complete the *S. agalactiae* HRC genome. Besides, we detected MinION reads consisting of low-complexity sequences compatible with artefacts of the technology but that did not affect the assembly.

## Supporting information

## Acknowledgments

Funded by Instituto de Salud Carlos III (PI14/00844; PI17/00610; FI17/00177) and Ministerio de Ciencia, Innovación y Universidades (RTC-2017-6471-1; MINECO/AEI/FEDER, UE) co-financed by the European Regional Development Funds, “A way of making Europe” from the European Union; and by the agreement OA17/008 with Instituto Tecnológico y de Energías Renovables (ITER) to strengthen scientific and technological education, training, research, development and innovation in Genomics, Personalized Medicine and Biotechnology.

## Authors’ contributions

T.H.B.: performed experiments, data analysis, interpretation, and manuscript drafting; H.R.P., A.D.U., R.G.M., J.L.S.: performed experiments and data analysis; F.L.D.: performed experiments, study conception, interpretation, revising the manuscript critically; C.F.: Study conception and design, data analysis, interpretation, revising the manuscript critically and conceived the project.

The authors declare that there are no competing interests.

## Data availability

All raw reads from MinION and Illumina are available from the SRA database (accession number(s) SRP141332, SRP141319).

