## Supplementary material for "Shallow MinION sequencing to assist *de novo* assembly of the *Streptococcus agalactiae* genome"

| **S1 Table.** Retrieved BLAST results for the MinION reads that did not align to the assembled *S. agalactiae* HRC genome. | | | | |
| --- | --- | --- | --- | --- |
| **Length (bp)** | **Alignment*** | **ID**** | **Query coverage** | **Identity** |
| 623 | Synthetic construct clone pRedCas9 | MF280109.1 | 99% | 91% |
|  | Red recombinase plasmid pKD46 | MF287367.1 | 99% | 91% |
|  | *E. coli* plasmid pCAGO | KY563785.1 | 99% | 91% |
|  | *E. coli* strain tolC | CP018801.1 | 99% | 91% |
| 5500 | *E. coli* strain tolC- | CP018801.1 | 98% | 90% |
|  | *E. coli* isolate 108 genome assembly | LT615379.1 | 98% | 90% |
|  | *E. coli* isolate 103 genome assembly | LT615378.1 | 98% | 90% |
|  | *E. coli* isolate 106 genome assembly | LT615377.1 | 98% | 90% |
| 6588 | *Shigella sp*. PAMC 28760 | CP014768.1 | 98% | 90% |
|  | *E. coli* strain C43(DE3) | CP011938.1 | 98% | 90% |
|  | *E. coli* strain C41(DE3) | CP010585.1 | 98% | 90% |
|  | *E. coli* BL21(DE3) | AM946981.2 | 98% | 90% |
| *Only top-four BLAST records are indicated.  **NCBI reference identifier. | |  |  |  |

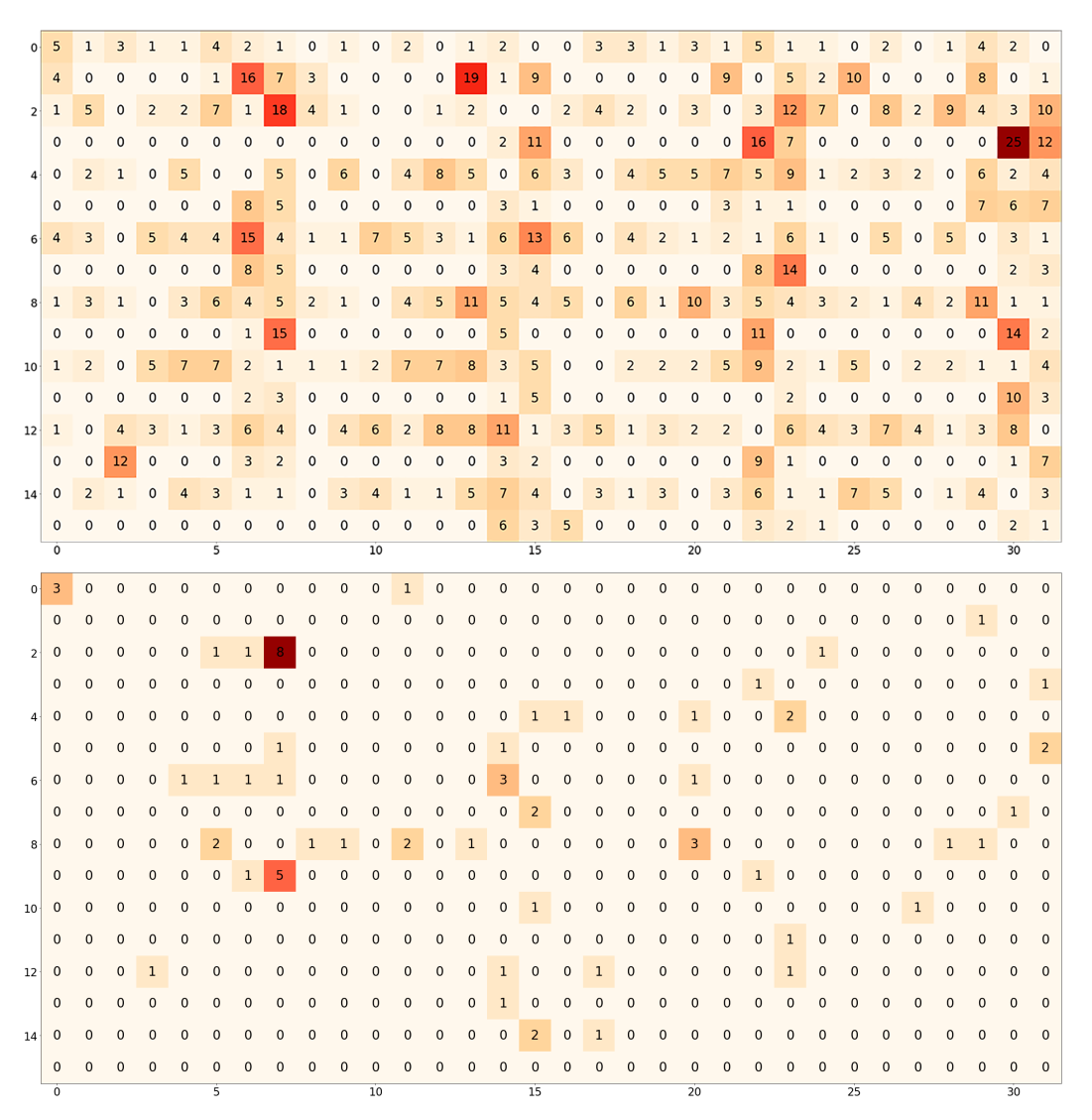

**S1 Fig.** Heatmap representing the number of reads per pore generated during the MinION experiment showing the 276 active pores that generated the 1,171 reads (top), and the 45 active pores involved in generating the reads that did not align (bottom). The gradient of colors indicates the number of reads.
